## Supplementary figures and tables for "Novel insights into the genetic architecture and mechanisms of host/microbiome interactions from a multi-cohort analysis of outbred laboratory rats"

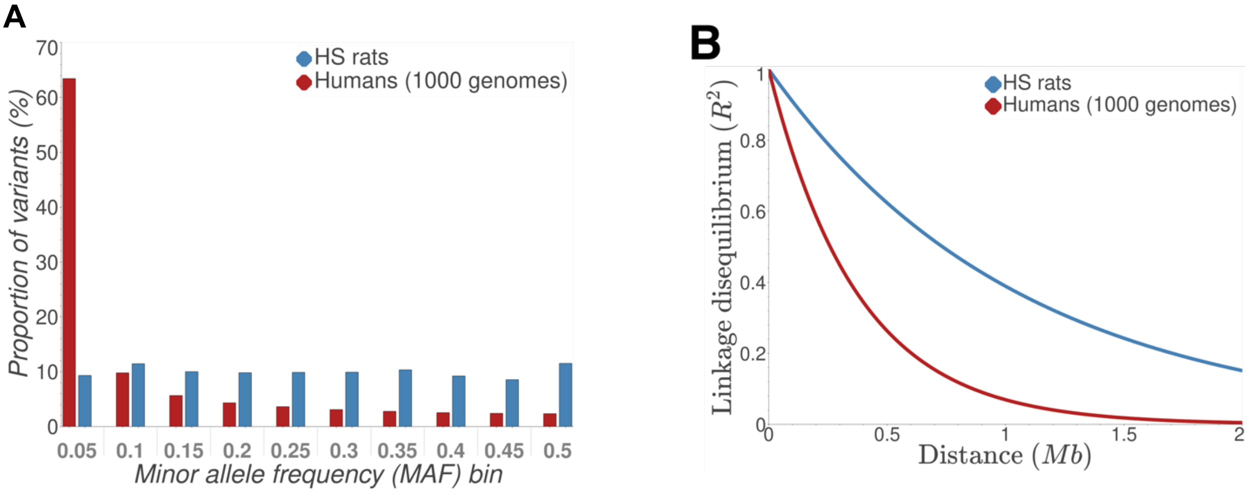


**Supplementary Figure 1 Genetic characteristics of the HS population favourable for GWAS.** (A) Comparison of the minor allele frequency (MAF) of genetic variants segregating in the HS rats and in humans. (B) Decay of linkage disequilibrium between neighbouring variants in the HS rats and in humans.

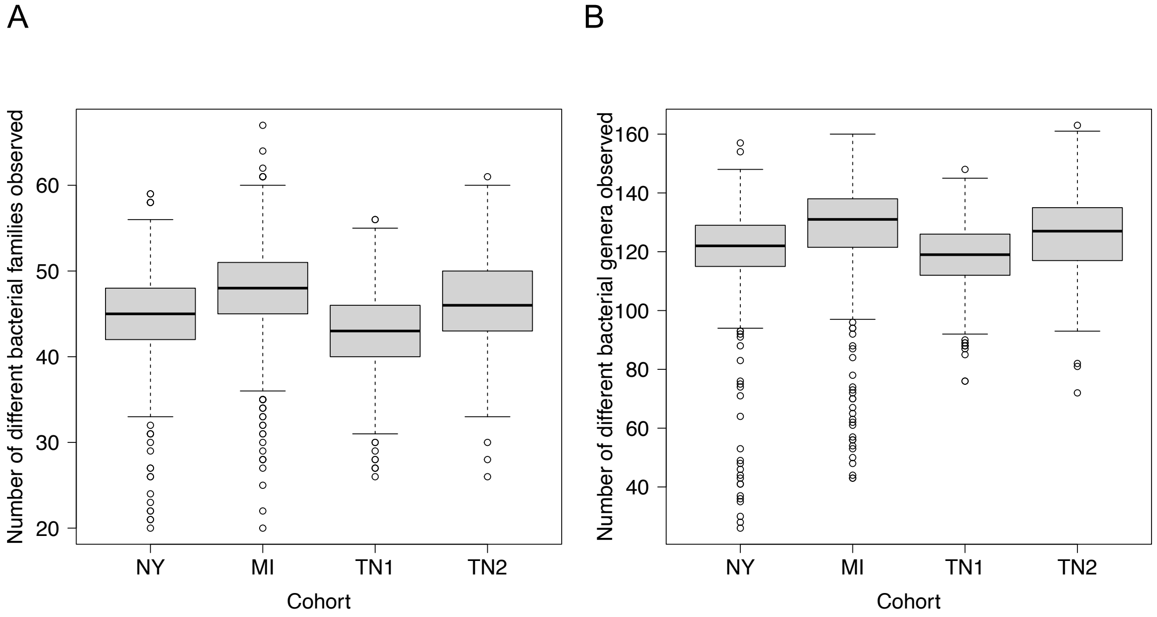


**Supplementary Figure 2 Within-sample (alpha) microbiome diversity.** Number of different (A) families or (B) genera observed in a sample (i.e. richness) in the different cohorts. A rarefied count table was used for this analysis.

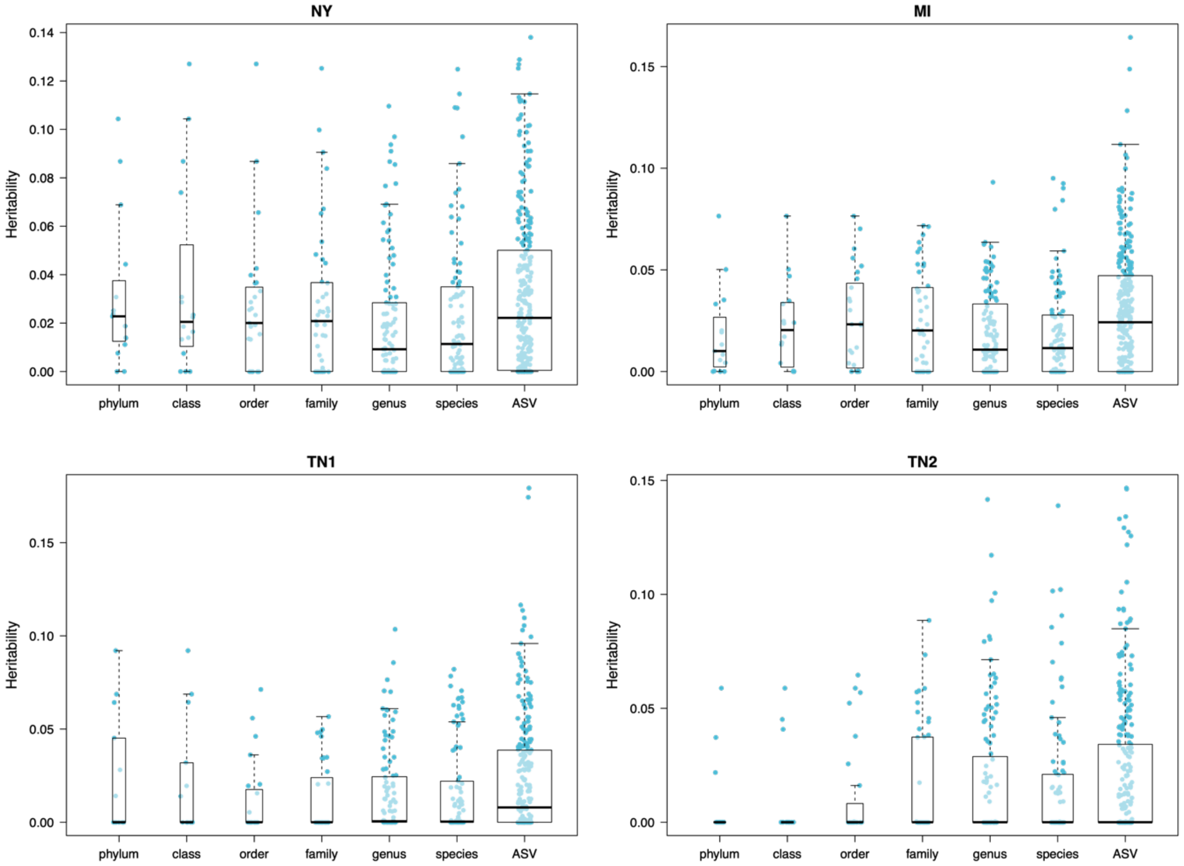


**Supplementary Figure 3 Comparison of heritability at different taxonomic levels.**

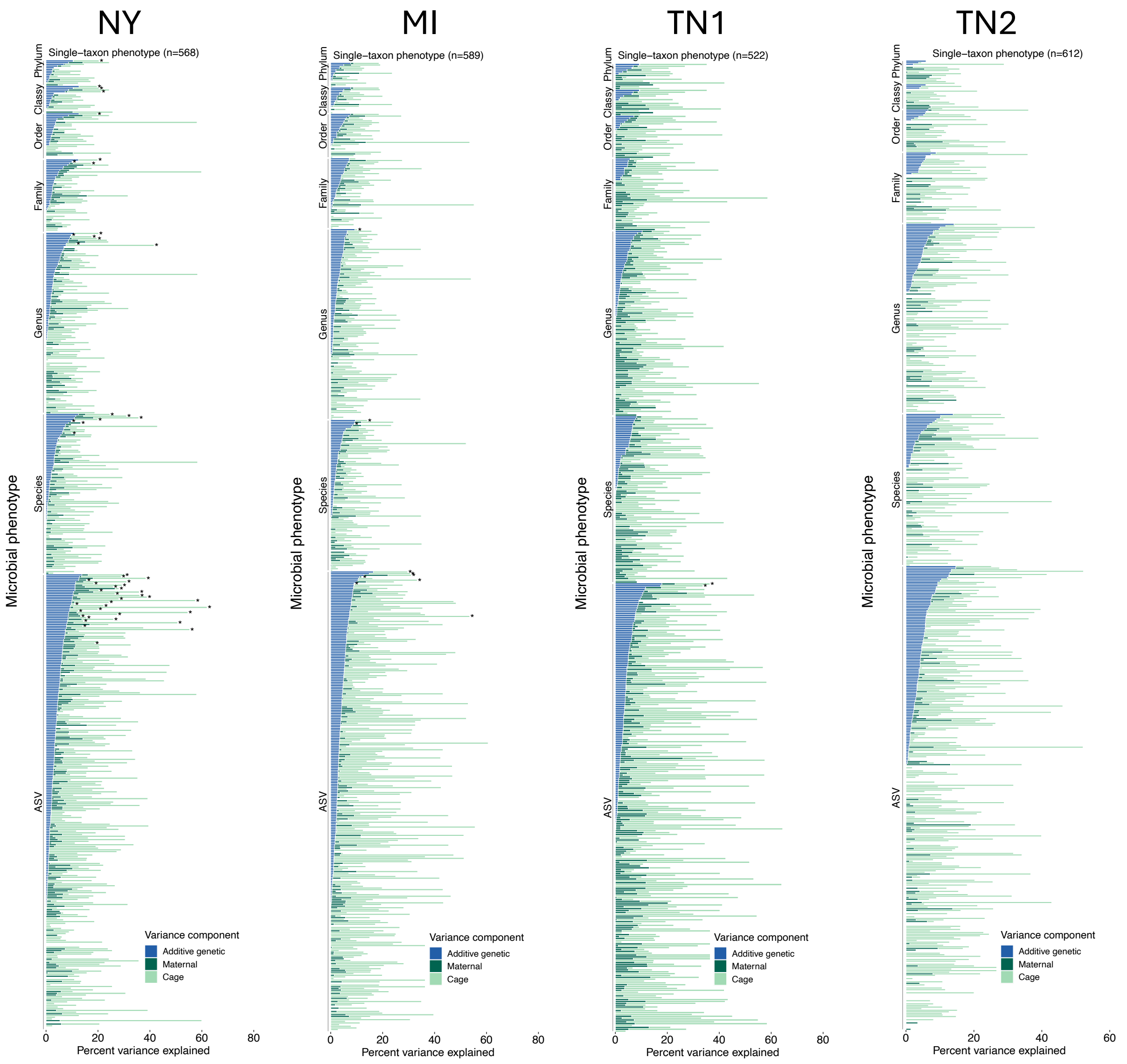


**Supplementary Figure 4 Decomposition of the variance of microbiome phenotypes.** Significant host genetic effects (cohort-wide FDR < 10%) are indicated with an asterisk. The code used to draw this figure is modified from the code provided by Grieneisen et al.[1].

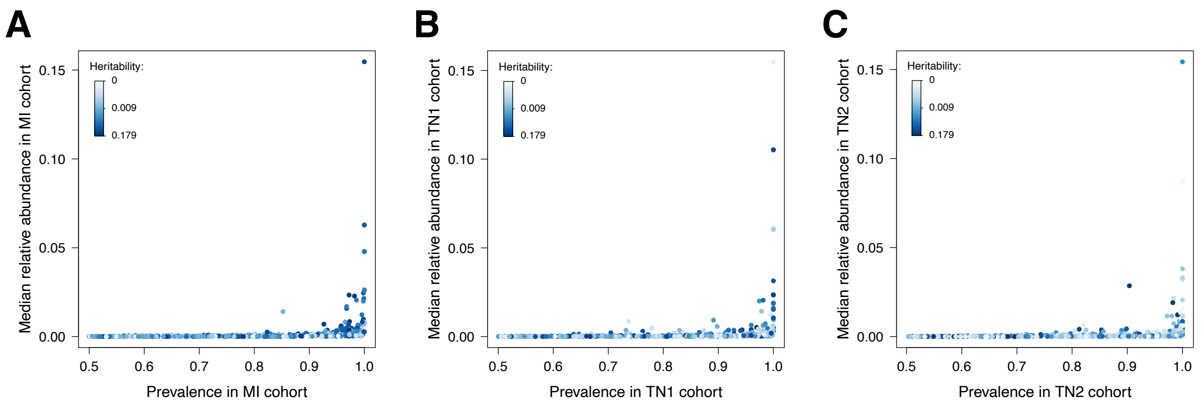


**Supplementary Figure 5 Relationship between prevalence, relative abundance and heritability for the microbiome phenotypes measured in the MI, TN1, TN2 cohorts**. The results for the NY cohorts are in main text (Fig. 3B).

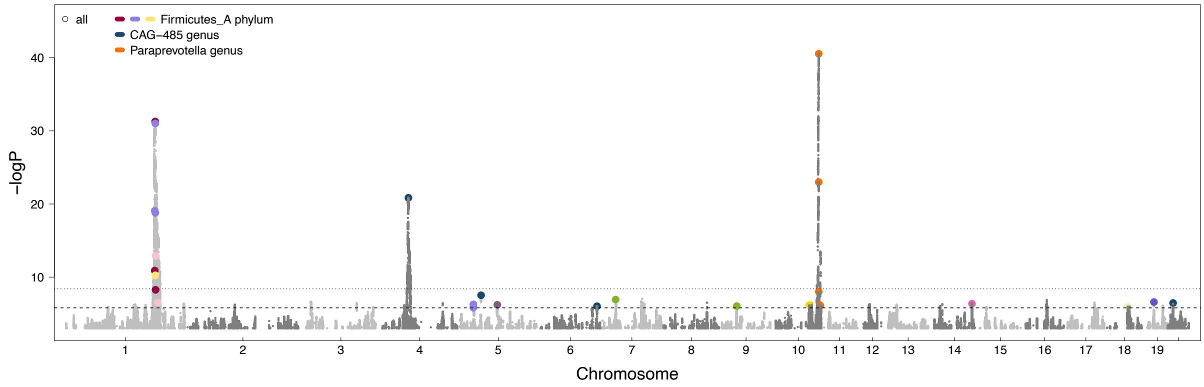


**Supplementary Figure 6 Microbiome-associated loci in the whole sample (N = 3,784).** This porcupine plot shows the association values for all 546 microbiome phenotypes with prevalence greater than 50% in the whole sample. The lower line (-logP = 5.8) reflects the genome-wide significance threshold for an individual trait, which accounts for the number of independent SNPs tested; the higher line (-logP=8.4) is the adjusted significance threshold, which in addition accounts for the number of independent microbiome phenotypes examined in each cohort. The colour of the dot refers to the genus affected; it is the same as in Fig.4. For that reason, genome-wide significant associations involving genera that did not have any genome-wide significant association in Fig. 4 are not higlighted by large coloured dots in this figure (but they are visible as grey peaks above the significance threshold).

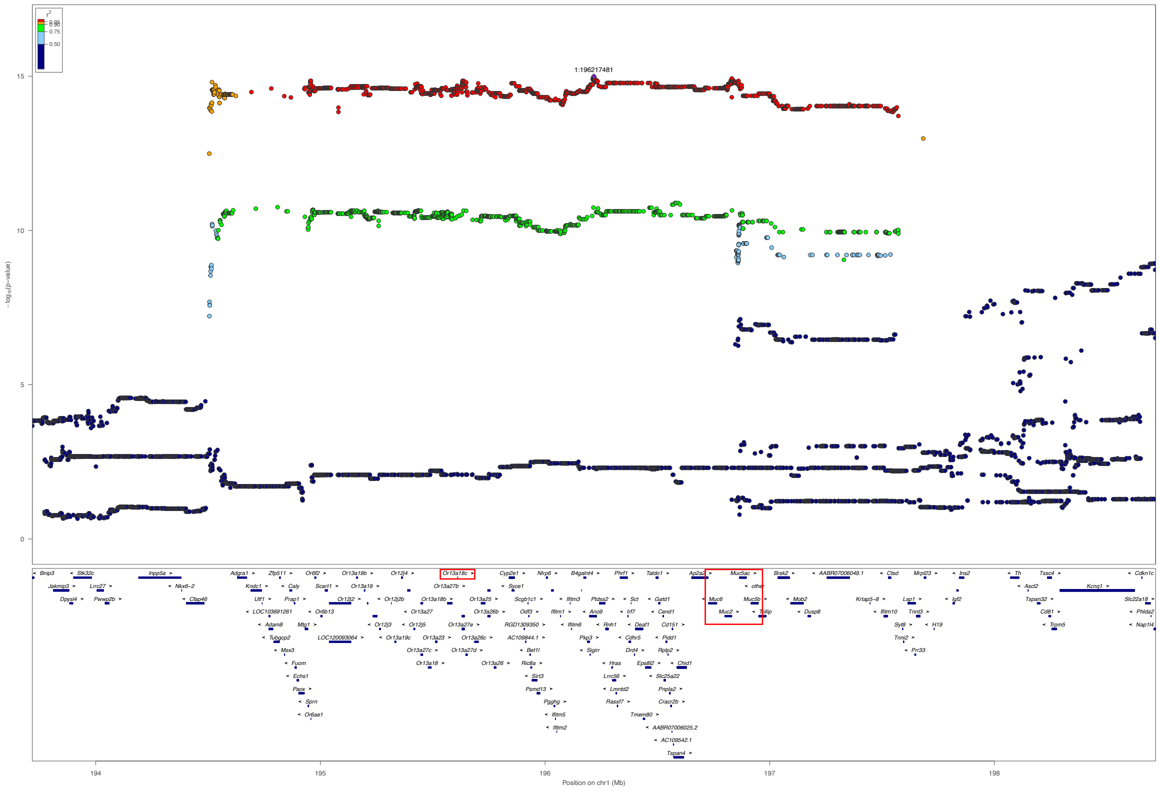


**Supplementary Figure 7 Association (LocusZoom) plot for ASV_3613 (order TANB77, family CAG-508) measured in the NY cohort, which is the phenotype with the most significant association with the chromosome 1 replicated locus.** The gene *Or13a18c* as well as the cluster of mucin-secreting genes (*Muc6, Muc2, Muc5ac and Muc5b*) are highlighted with red rectangles.

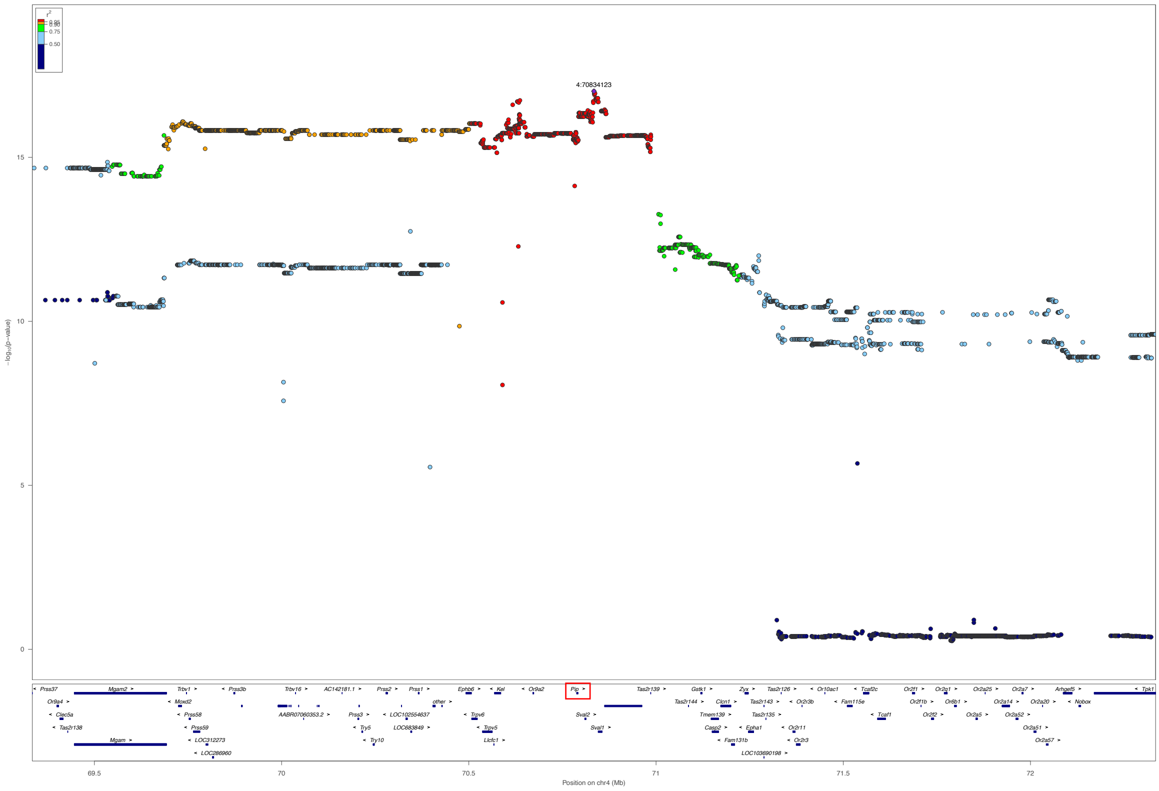


**Supplementary Figure 8 Association plot for ASV_18566 (family Muribaculaceae, genus *CAG-485*, species *002362485*) measured in the NY cohort, which is the phenotype with the most significant association with the chromosome 4 replicated locus.** The gene *Pip* is highlighted with a red rectangle.

A

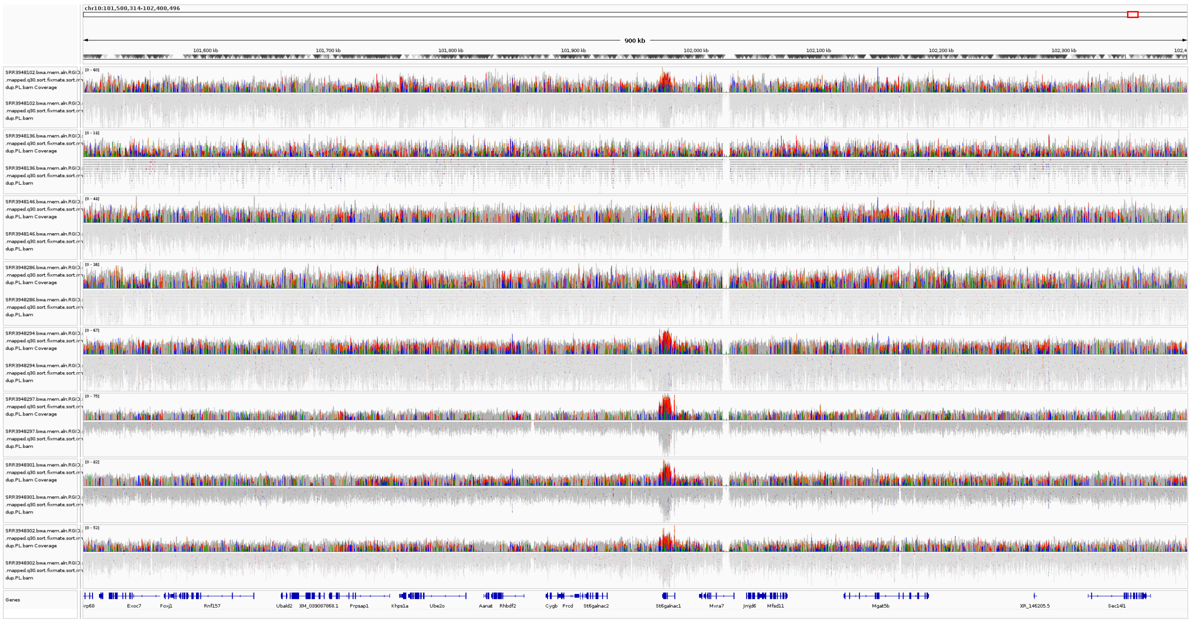


B

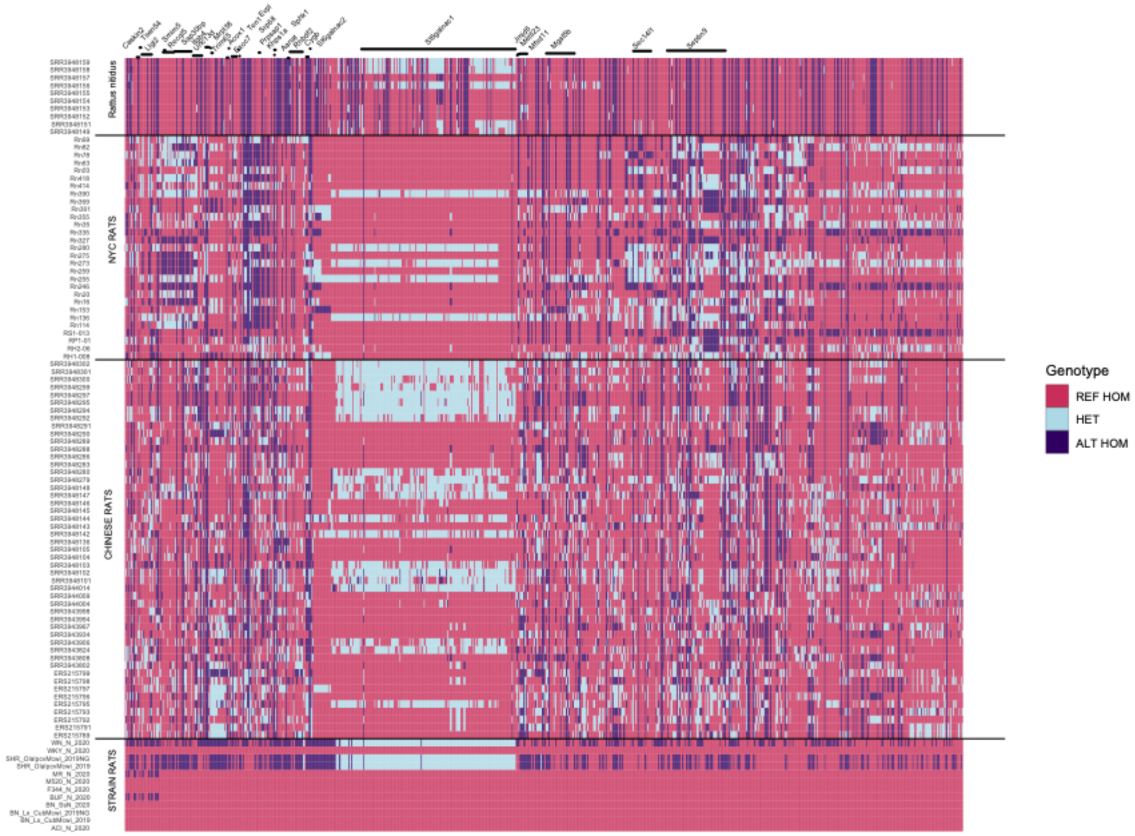


**Supplementary figure 9 Evidence that the copy number variant affecting *St6galnac1* and seen in the WN/N HS founder and outbred HS rats (Fig. 5B) also segregates in wild rats.** (A) Short read sequencing of wild rats captured in various locations in China and sequenced in Hiseq Ten X (data publicly available from Teng et al. 2017[2]). The region surrounding the *St6galnac1* gene is shown. The bump in sequencing depth over *St6galnac1* shows that the same partial duplication/triplication of the gene segregates in wild rats from China as in the HS. (B) Genotypes of the eight HS founders (including WN/N) and two SHR inbred strains (bottom), wild rats from China (middle) and wild rats from New York City (VCF available from Harpak et al. 2020[3]) (top) showing a similar pattern of apparent variants accumulation in the region corresponding to the copy number variant. This apparent accumulation of variants is due to the presence of paralogous sequences, and results in apparent heterozygosity of the inbred strains.

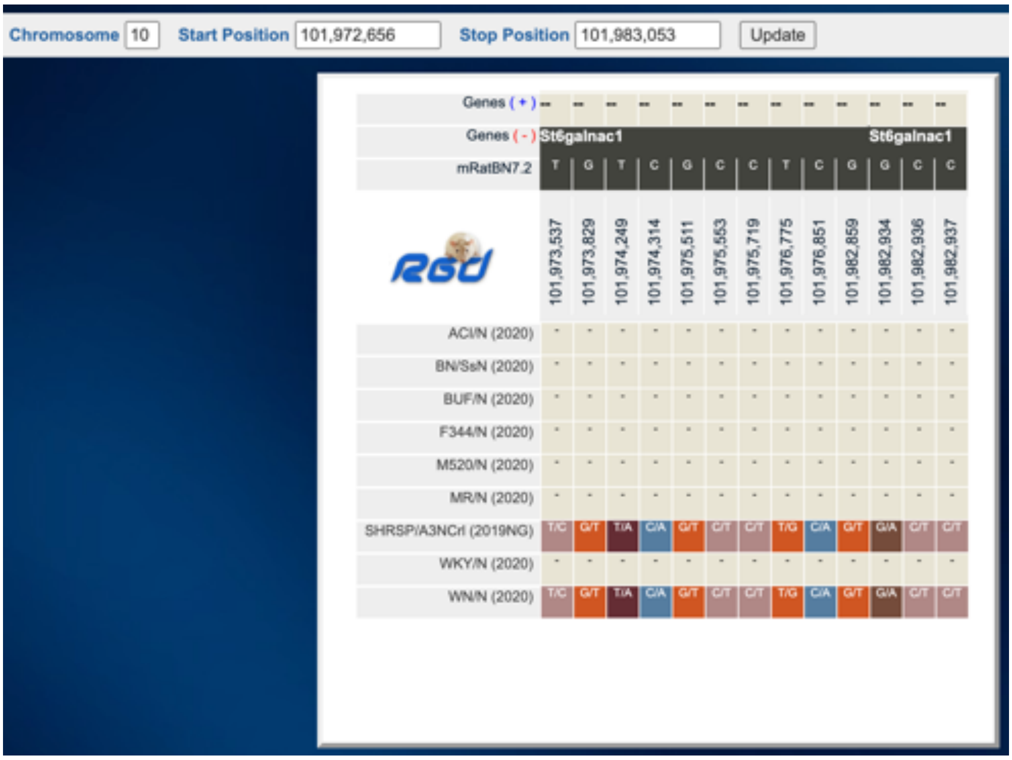


**Supplementary figure 10 Screenshot from the Variant Visualizer of the Rat Genome Database[4] showing all variants identified in the eight HS founders and in the strain SHRSP/A3N (not an HS founder).** This figure shows that WN/N and SHRSP/A3N are identical by descent in this region.

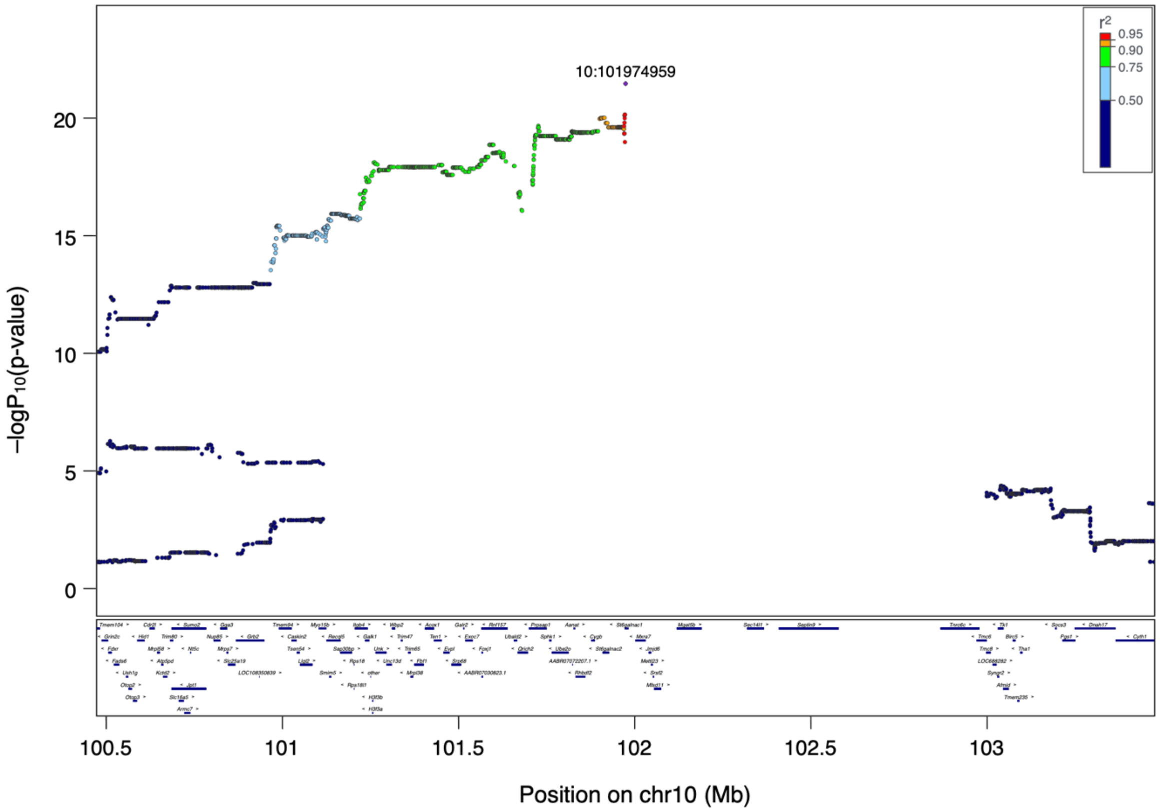


**Supplementary figure 11. Local association plot for ASV_5163 (genus *Paraprevotella*) measured in the NY cohort on chromosome 10** (plot for zoomed in region is shown in Fig. 5A). The gap between 102-103Mb is not a technical issue: variants are detected between the eight inbred founders of the HS population but not in the outbred population. In that region, seven of the founders are identical by descent and the eighth founder (WN/N) is different. The WN/N haplotype was lost at some point during outbreeding, before reaching the generations included in this study.

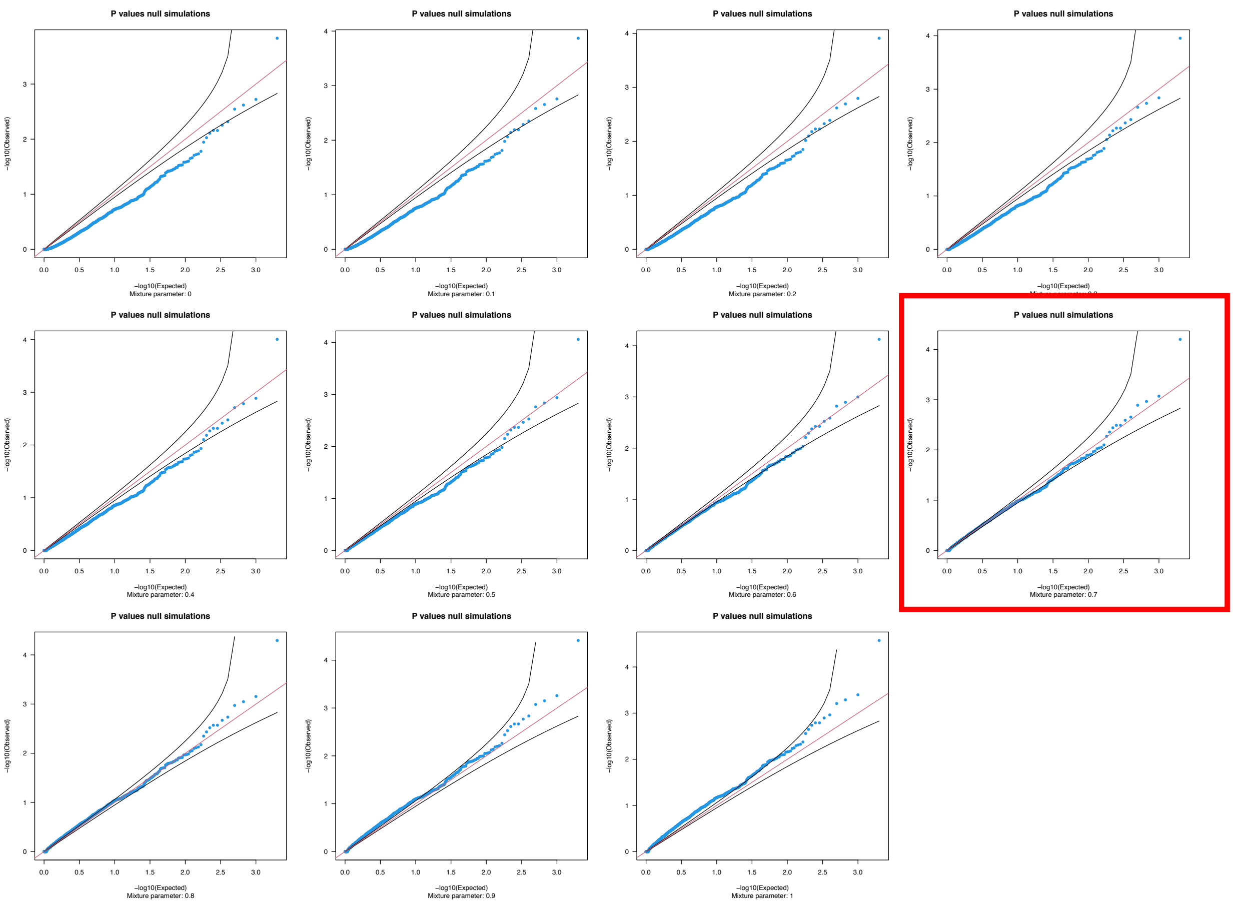


**Supplementary Figure 12 Calibration of Mic-IGE *p*-values using null simulations.** We chose 0.7 as the most appropriate parameter for the mixture of chi-square distributions with degrees of freedom 1 and 2 used to obtain Mic-IGE *p*-values. The resulting *p*-values obtained for null simulations are calibrated.

See Methods section “Evaluation of the calibration of *p*-values using null simulations” for full details.

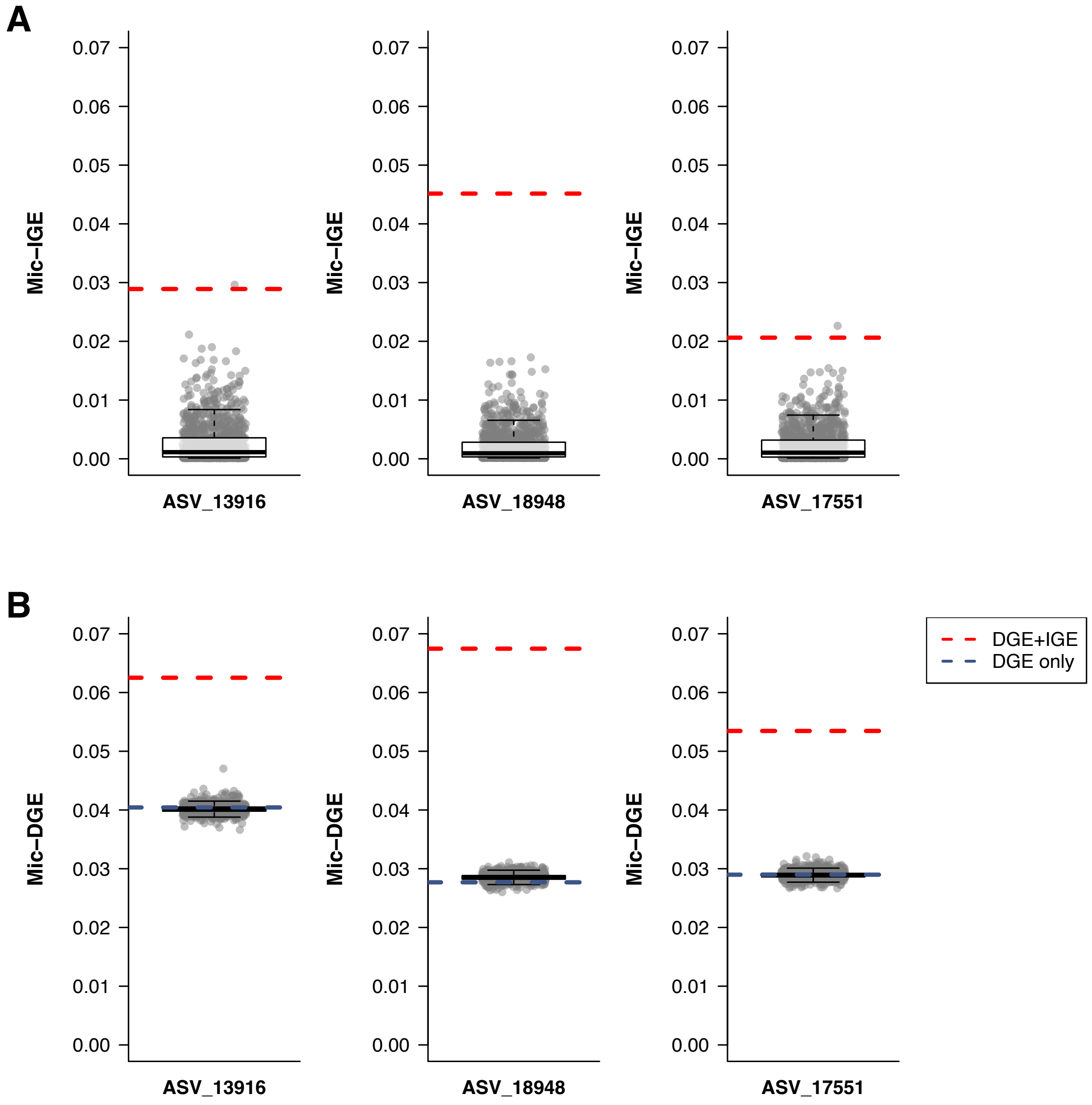


**Supplementary Figure 13 Variance explained by Mic-IGE (A) and Mic-DGE (B) after permutation of cage mates' assignments.** The cage assignments, used to account for cage effects, were not permuted.

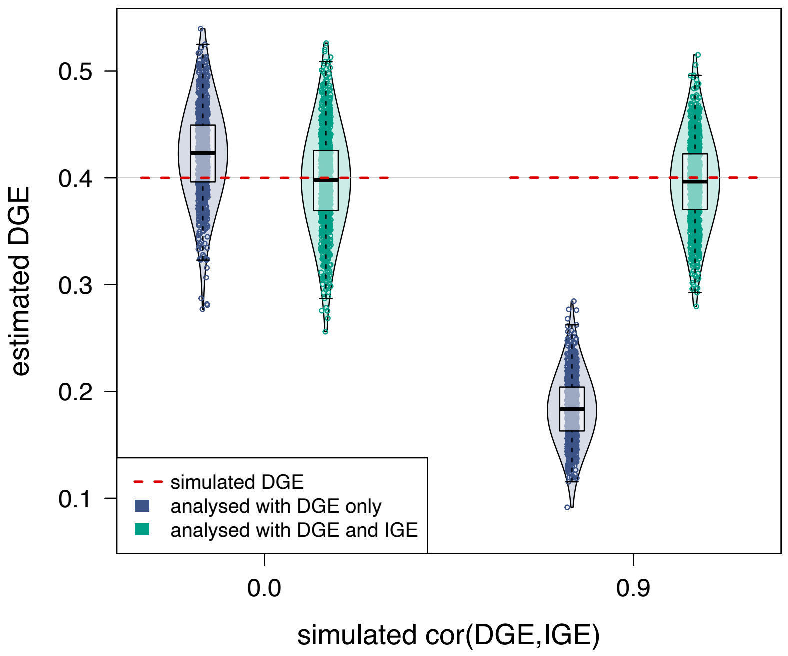


**Supplementary Figure 14 Simulation study showing that DGE estimates are biased (underestimated) when IGE are at play but not modelled.** Phenotypes were simulated for the rats in the MI cohort (see Figure 6D for simulations for the rats in the NY cohort, and Methods for details on the simulations).

| | Cohort | | | |
|---|---|---|---|---|
| | **NY** | **MI** | **TN1** | **TN2** |
| **Facility rats were born in** | MCW | MCW | TN | MCW |
| **Facility rats were reared in, after weaning and shipment** | Univ. at Buffalo, NY | Univ. of Michigan, MI | Univ. of Tennessee, TN | Univ. of Tennessee, TN |
| **Number of rats per cage** | 2 | 3 | 1-4 | 2 |
| **Sexes of co-housed rats** | same-sex | same-sex | same-sex | one male, one female (breeding) |
| **Relatedness of co-housed rats** | unrelated | unrelated | littermates | unrelated |
| **Specific pathogen free facility** | yes | no | yes | yes |
| **Type of cages** | conventional | conventional | conventional & isolators | conventional & isolators |
| **Type of diet** | chow | chow | chow | chow |
| **Exact diet** | Teklad 18% Protein Rodent Diet (Envigo) | Picolab Laboratory Rodent Diet (LabDiet) | Teklad Irradiated LM-485 Mouse/Rat Diet (Envigo) | Teklad Irradiated LM-485 Mouse/Rat Diet (Envigo) |
| **Behavioural phenotyping*** | locomotion, cocaine conditioning and behavioural regulations | cocaine sensitization and Pavlovian cue-reactivity | socially-acquired nicotine self-administration | none |
| **Drugs taken for phenotyping** | cocaine (intraperitoneal) | cocaine (intraperitoneal) | nicotine (intravascular) | none |
| **Age when ceca collected (months)** | 6.5 | 3 | 2 | 6 |
| **Overnight fasting prior to sacrifice** | yes | yes | no | yes |
| **Chemical used for euthanisia method** | isoflurane (inhalation) | none | isoflurane (inhalation) | isoflurane (inhalation) |
| **Physiological phenotyping (just before/after euthanasia)** | fasting glucose, adiposity | fasting glucose, adiposity | adiposity | fasting glucose, adiposity |
| **Cohort size (post filtering)** | 1167 | 1112 | 950 | 555 |

* for a detailed description of the protocols used, see ratgenes.org, tab NIDA Research

**Supplementary Table 1 Between-cohort environmental and experimental differences.**

| ASV or taxon | Cohort | Chr. | Position | -logP | Taxonomy |
|---|---|---|---|---|---|
| f__CAG-74 | MI | 1 | 196,695,784 | 9.9** | o__Christensenellales;f__CAG-74 |
| ASV_59945 | MI | 1 | 196,831,050 | 7.6* | o__Christensenellales;f__CAG-74;g__Limiplasma |
| g__Limiplasma | MI | 1 | 196,831,050 | 6.6* | o__Christensenellales;f__CAG-74;g__Limiplasma |
| s__Limiplasma_merdipullorum | MI | 1 | 196,831,050 | 6.7* | o__Christensenellales;f__CAG-74;g__Limiplasma |
| ASV_63700 | MI | 1 | 195,080,343 | 9.0* | o__Christensenellales;f__CAG-74;g__Onthenecus |
| g__Onthenecus | MI | 1 | 195,080,344 | 7.2* | o__Christensenellales;f__CAG-74;g__Onthenecus |
| ASV_70101 | MI | 1 | 196,498,032 | 5.8* | o__Oscillospirales;f__Oscillospiraceae_88309;g__Limivicinus |
| ASV_2972 | MI | 1 | 195,080,344 | 6.5* | o__TANB77;f__CAG-508;g__CAG-269 |
| ASV_26581 | NY | 1 | 197,683,398 | 8.6** | o__TANB77;f__CAG-508;g__CAG-273 |
| ASV_26581 | TN_breeder | 1 | 196,878,658 | 6.0* | o__TANB77;f__CAG-508;g__CAG-273 |
| ASV_29320 | TN_behavior | 1 | 196,498,112 | 12.5** | o__TANB77;f__CAG-508;g__CAG-793 |
| ASV_29320 | MI | 1 | 195,701,712 | 6.3* | o__TANB77;f__CAG-508;g__CAG-793 |
| g__CAG-793 | NY | 1 | 195,629,919 | 7.5* | o__TANB77;f__CAG-508;g__CAG-793 |
| s__CAG-793_sp000433915 | NY | 1 | 195,629,919 | 9.5* | o__TANB77;f__CAG-508;g__CAG-793 |
| **ASV_3613** | **NY** | **1** | **196,217,481** | **15.0**** | **o__TANB77;f__CAG-508;g__UMGS1994** |
| ASV_3613 | MI | 1 | 197,534,412 | 10.6** | o__TANB77;f__CAG-508;g__UMGS1994 |
| ASV_3613 | TN_breeder | 1 | 197,572,275 | 6.9* | o__TANB77;f__CAG-508;g__UMGS1994 |
| g__UMGS1994 | TN_behavior | 1 | 196,916,401 | 6.9* | o__TANB77;f__CAG-508;g__UMGS1994 |
| g__UMGS1994 | MI | 1 | 197,208,428 | 6.4* | o__TANB77;f__CAG-508;g__UMGS1994 |
| s__UMGS1994_sp900553945 | MI | 1 | 197,208,428 | 7.0* | o__TANB77;f__CAG-508;g__UMGS1994 |
| s__UMGS1994_sp900553945 | TN_behavior | 1 | 196,916,401 | 6.6* | o__TANB77;f__CAG-508;g__UMGS1994 |

**Supplementary Table 2 Significant associations at the chromosome 1 pleiotropic and replicated locus.** Genome-wide significance (-logP > 5.8) is shown with an asterisk (*); Bonferroni-adjusted significance (-logP > 8.4) with two (**). Only associations in strong LD with the lead SNP (shown in bold) are included in the table. All these ASVs and taxa belong to the Firmicutes_A phylum (GTDB taxonomy). Row colours correspond to the colours in the porcupine plot (Fig. 4). The first row (white) refers to a microbiome phenotype that does not appear in the porcupine plot because it is only annotated at the family level, not at the genus level.

| ASV or taxon | Cohort | Chr. | Position | -logP | Taxonomy |
|---|---|---|---|---|---|
| **ASV_18566** | **NY** | **4** | **70,834,123** | **17.0**** | **o__Bacteroidales;f__Muribaculaceae;g__CAG-485** |
| ASV_18566 | MI | 4 | 70,349,586 | 7.0* | o__Bacteroidales;f__Muribaculaceae;g__CAG-485 |
| ASV_18566 | TN_behavior | 4 | 70,473,002 | 5.9* | o__Bacteroidales;f__Muribaculaceae;g__CAG-485 |

**Supplementary Table 3 Significant associations at the chromosome 4 replicated locus.** Genome-wide significance (-logP > 5.8) is shown with an asterisk (*); Bonferroni-adjusted significance (-logP > 8.4) with two (**). Only associations in strong LD with the lead SNP (shown in bold) are included in the table. Row colours correspond to the colours in the porcupine plot (Fig. 4).

| ASV or taxon | Cohort | Chr. | Position | -logP | Taxonomy | Direction of effect |
|---|---|---|---|---|---|---|
| ASV_5096 | MI | 10 | 101,922,691 | 10.4** | o__Bacteroidales;f__Bacteroidaceae;g__Paraprevotella | WN/N decreasing |
| **ASV_5163** | **NY** | **10** | **101,974,959** | **21.5**** | **o__Bacteroidales;f__Bacteroidaceae;g__Paraprevotella** | **WN/N decreasing** |
| ASV_5163 | TN_behavior | 10 | 101,972,949 | 8.5** | o__Bacteroidales;f__Bacteroidaceae;g__Paraprevotella | WN/N decreasing |
| ASV_5163 | TN_breeder | 10 | 101,769,054 | 4.2 | o__Bacteroidales;f__Bacteroidaceae;g__Paraprevotella | WN/N decreasing |
| g__Paraprevotella | NY | 10 | 101,727,325 | 12.5** | o__Bacteroidales;f__Bacteroidaceae;g__Paraprevotella | WN/N decreasing |
| g__Paraprevotella | MI | 10 | 101,938,835 | 10.3** | o__Bacteroidales;f__Bacteroidaceae;g__Paraprevotella | WN/N decreasing |
| g__Paraprevotella | TN_breeder | 10 | 101,905,255 | 4.1 | o__Bacteroidales;f__Bacteroidaceae;g__Paraprevotella | WN/N decreasing |
| ASV_2821 | MI | 10 | 101,974,959 | 4.0 | o__Verrucomicrobiales;f__Akkermansiaceae;g__Akkermansia | WN/N decreasing |
| ASV_17008 | MI | 10 | 101,972,949 | 4.5 | o__Bacteroidales;f__Muribaculaceae;g__Muribaculum | WN/N increasing |

**Supplementary Table 4 Significant associations and suggestive (logP > 4) at the chromosome 10 replicated locus.** Genome-wide significance (-logP > 5.8) is shown with an asterisk (*); Bonferroni-adjusted significance (-logP > 8.4) with two (**). Only associations in strong LD with the lead SNP (shown in bold) are included in the table. Row colours correspond to the colours in the porcupine plot (Fig. 3).
